# Deciphering the molecular mechanism associated with the interaction of TULP3 and Histone deacetylases, SIRT1 and SIRT2

**DOI:** 10.1101/2024.12.23.630205

**Authors:** Ramu Manjula, Gopinatha Krishnappa, Vikeraman Sriram, Padmanabhan Balasundaram

**Affiliations:** Department of Biophysics, National Institute of Mental Health and Neurosciences (NIMHANS), Hosur Road, Bengaluru – 560029, India

**Keywords:** TULP3, Sirtuins, protein-protein interactions, biophysical assays, biochemical assays, deacetylate activity

## Abstract

Tubby-like protein-3 (TULP3), a member of the tubby family, is recognized for its role in ciliary trafficking, linking membrane-associated proteins to the intraflagellar transport complex A (IFT-A). Mammalian tubby family proteins (TULP1-4), featuring a C-terminal tubby domain, play crucial roles in intracellular transport, signaling, cell differentiation, and motility. TULP3 is also associated with renal and hepatic fibrocystic disease and liver, kidney, and heart degeneration. However, the modulation of TULP3 function and its interacting partners in the cell still need to be understood. Recent studies have indicated interactions between TULP3 and various proteins, including acetylating and deacetylating enzymes. In this study, we investigate the interaction of TULP3 with Sirtuins, SIRT1, and SIRT2 using biochemical and biophysical techniques. Our findings revealed that both the N- and C-terminal domains of TULP3 are necessary for the interaction with SIRT1 and SIRT2. Furthermore, we have identified that TULP3 is not a deacetylation substrate for SIRT1.

## 1. Introduction

Tubby-like proteins (TULPs), characterized by their highly conserved tubby domain, are present in both plant and animal kingdoms. Among the four TULPs (TUB and TULP1–3), they are predominantly localized in the nervous tissue [1–7], and they exhibit dual localization within the nucleus and plasma membrane [1]. While TULP1-4 is expressed in the retina [2,3], TUB and TULP3 demonstrate expression in the central nervous system and various tissues, indicating a broader tissue expression pattern than TULP1 and TULP2. High RNA levels for TULP3 have been detected in the spinal cord, ovaries, thyroid, and testis [3]. Structural studies have revealed that TULP proteins act as transcription factors, featuring a C-terminal tubby domain capable of binding to double-stranded DNA and selectively interacting with membrane phosphoinositides [4,5]. The N-terminal region resembles the transcriptional activation region of known transcription factors [4]. Tubby proteins function through signal transduction pathways through G-protein coupled receptors (GPCRs) which translocate the tubby from the plasma membrane to the nucleus [1]. TULP3 stands out as the sole member in the family implicated in embryonic development, with mutations in the TULP3 gene leading to abnormalities in neuronal tube development in mice [6,7]. This is attributed to TULP3’s role in inhibiting Hedgehog signaling, thereby regulating the patterning of mouse embryos [7]. Notably, TULP3 mutations have also been associated with renal and hepatic fibrocystic disease and hypertrophic cardiomyopathy [8,9].

Though there are primary structural clues, potential targets of tubby are not known completely, and least is known about the signaling pathways and molecules regulating the function of TULP proteins. The N-terminus conserved helix of TULP3 binds to the intraflagellar transport-A (IFT-A) complex to promote ciliary G-protein coupled receptor (GPCR) localization [5,10–12]. At lower concentrations of phosphatidylinositol 4, 5-bisphosphate (PIP2), TULP3 interacts with the IFT-A complex to transport 16 different classes of GPCRs (melanin-concentrating hormone receptors, neuropeptide Y receptors, and GPR161) and other integral membrane proteins (Polycistin 1/2 complex) [11,13,14]. A study conducted has successfully identified novel interactors that play a crucial role in regulating the activity and stability of the TULP3 protein in cells. The research demonstrates that acetylation by p300 and deacetylation modifications by Histone deacetylases (HDACs) are instrumental in regulating TULP3 ubiquitination [15]. In this study, the author has validated the TULP3 interaction with deacetylase HDAC1 and SIRT1, a ubiquitin ligase RAD18, and protein phosphatase 6 regulatory subunit3 (PP6R3). Interestingly, another study by has further supported the interaction between TULP3 and SIRT1. This interaction is implicated in critical cellular processes such as DNA damage repair and fibrosis [9]. These findings shed light on the intricate regulatory network governing TULP3 protein dynamics in cellular functions.

Sirtuins, anti-aging proteins, are conserved from bacteria to humans with similar structures and catalytic functions. Sirtuins, which belong to the class III histone deacetylases (HDACs), catalyze NAD+-dependent deacetylation [16] and engage in various activities such as ribosyltransferase [17], demalonylase, and desuccinylase processes. SIRT1 and SIRT2 belong to the sirtuins family. Studies show sirtuins slow aging in mammals, extending lifespan through caloric restriction (CR) and supporting cell survival during CR [18,19]. They demonstrate a pro-survival role by inhibiting apoptosis and fostering cell survival [17]. SIRT1 is linked to neurodegenerative disorders, cancers, anti-inflammatory effects, autophagy, cell senescence, and aging [20,21]. Meanwhile, SIRT2’s roles include apoptosis, necroptosis, autophagy, neuroinflammation, immune response, and tumorigenesis regulation with tumor-promoting and tumor-suppressing functions [22,23].

Further exploration is required to comprehensively understand the interaction between SIRT1/2 and TULP3 and the regulatory mechanisms influencing cellular functions upon their interaction.

In our study, we employed modified constructs of TULP3, SIRT1, and SIRT2 to investigate their interactions using biochemical and biophysical techniques, including pull-down assay, surface plasmon resonance (SPR), and Microscale thermophoresis (MST). The binding affinities of specific TULP3 constructs were individually assessed against SIRT1 and SIRT2 proteins to identify the interacting domains. Understanding the intricate details of these molecular interactions is crucial for unraveling the cellular functions of TULP3 and its role in modulating sirtuin activity. These findings pave the way for future investigations into the broader cellular pathways regulated by the TULP3-Sirtuin axis and its relevance in physiological and pathological contexts.

## 2. Materials and Methods

### 2.1. Bacterial strains

All bacterial strains used in this study were derived from *E. coli* (*Escherichia coli*) K-12. The cloning host, DH5α, was used for cloning and expression hosts, BL21 (DE3) codon plus (Invitrogen), and Rosetta 2 (Merck) strains were used for overexpression of recombinant proteins.

### 2.2. Cloning of TULP3, SIRT1 and SIRT2

The hTULP3 (cDNA-HsCD00329661) plasmid was sourced from the DNASU plasmid repository at Arizona State University. PCR products for the N-terminal hTULP3 (1-148 aa, construct-1) and deleted hTULP3 (60-442 aa, construct-2) were generated using specific primers. For construct-1, the forward primer was 5’-GATGCGGGATCCATGGAGGCTTCGCGCTG-3’, and the reverse primer was 5’-GATGCGCTCGAGTCACTGGGATATTCCATCAGTCTCC-3’. For construct-2, the forward primer was 5’-GATGCGGGATCCGCAAAGCCAAGGGCCAG-3’, and the reverse primer was 5’-GATGCGCTCGAGTCACAATTCACACGCCAGC-3’. These constructs were inserted between the BamHI and XhoI restriction sites into the pGEX-6P1 bacterial expression vector, featuring a GST tag at the N-terminal region. Sequencing confirmed the integrity of the pGEX-6P1-TULP3 constructs.

Additionally, PCR products of hSIRT1 (114-510 aa) was amplified by using primers ( forward primer: 5’-ATGCCATGGcaATTGGGTACCGAG-3’ and reverse primer: 5’-CCGCTCGAGCTATCGTGGAGGTTTTTCAGTAATTTC-3’) and hSIRT2 (36-356 aa) was amplified by using primers (forward primer: 5’-ATGCCATGGCAGGAGAAGCAGACATGGAC-3’ and reverse primer: 5’-ACGTCGCTCGAGTTACGACTGGGCATCTATGCTG-3’) and amplified PCR products were inserted between the NdeI and XhoI restriction sites into the pET28a bacterial expression vector, which carries a hexa Histidine-tag at the N-terminal region. Sequencing confirmed the correctness of the pET28a, hSIRT1, and hSIRT2 clones.

### 2.3. Expression and protein purification of TULP3, hSIRT1 and hSIRT2

The protein expression conditions for TULP3 constructs were standardized in the Rosetta (DE3) bacterial expression system, induced with 0.5 mM IPTG, and incubated at 18°C for 16 hrs post induction. Protein was purified from the pellet stored at −80 °C. Pellet was allowed to thaw on ice for 30 min, and the lysis buffer (50 mM Tris, pH 8.0, 300 mM NaCl, 5mM DTT, 1mM PMSF, and 10 % glycerol) containing lysozyme and DNaseI was added to dissolve pellet. Once the pellet was dissolved, the lysate was on ice for 15 min. Liquid nitrogen was used to freeze the lysate and then allowed to thaw under tap water. Freeze-thaw was repeated twice, and the lysate was taken for sonication using VibraCell (30 % Amplitude, 3 sec on and 5 sec off). The cell lysate was centrifuged by ultracentrifugation at 20,000 rpm for 30 min at 4 °C. The TULP3 supernatant was collected and loaded into a 5 ml HiTrap HP, GST affinity column (GE Healthcare). The fractions containing pure TULP3 proteins were pooled and desalted against 20 mM Tris pH 8.0, 300 mM NaCl, 5 mM DTT, 1 mM PMSF, and 10 % glycerol, using Hiprep 23/10 desalting column (GE Healthcare). The purified proteins were taken for the size exclusion chromatography column (Superdex 75G 16/600, GE Healthcare) using the AKTA Prime purification system (GE Healthcare). The pure proteins were concentrated, aliquoted, and stored at −80°C for biochemical assays.

Sirtuin proteins (SIRT1 and SIRT2) were purified as described previously [24]. Briefly, proteins were expressed in Rosetta 2 cells using 0.5 mM IPTG at 16°C for 16 hours. After sonication, the soluble proteins were collected by ultracentrifugation and subjected to Ni-NTA affinity purification. Pure sirtuin fractions were desalted, and thrombin-digested proteins were further purified using size exclusion chromatography. The final purified proteins were concentrated, aliquoted, and stored at −80°C for biochemical analysis.

### 2.4. Pull-down assay

For pull-down experiments, SIRT1 and SIRT2, with an N-terminal His tag, functioned as the immobilized bait, with TULP3 acting as the prey. After expressing sirtuin and TULP3 constructs as previously described, a 10 ml cell pellet was utilized for the analysis. Before sonication, the cell pellet was dissolved in 400 µl lysis buffer (20 mM Tris, pH 8.0, and 150 mM NaCl). For one hour, SIRT1 and SIRT2 supernatants were incubated with 50 µl of Ni-NTA beads at 4°C. After this incubation period, the flow-through was collected, and the beads underwent three washes with 0.5 ml Tris-buffered saline (TBS). Following these washes, the prey supernatant was loaded onto the beads and allowed to bind to Sirtuins during a one-hour incubation at 4°C. Subsequently, the flow-through was collected, and the beads were washed three times with TBS buffer. The prey was then pulled down with the bait using 300mM imidazole. All collected samples were loaded onto SDS-PAGE and analyzed by Western blotting.

### 2.5. Western blot

Following gel electrophoresis, proteins were transferred to PVDF membranes using a BioRad Trans-Blot SD Semi-Dry Transfer Cell. PVDF membranes and blotting filter paper were pre-soaked in chilled Transfer buffer (0.025 M Tris, 0.192 M glycine, 20% v/v methanol). Gel equilibrated in Transfer buffer was transferred at 200 mA for 2 hours. PVDF membranes were immediately used for antibody probing. A 2-hour blocking in 3% w/v non-fat milk in PBST was followed by overnight rocking at 4 °C with primary antibodies (anti-His-Abcam, Rabbit polyclonal (Ab137839) and anti-TULP3-Abcam, Rabbit monoclonal (Ab138816)). After washing, secondary antibodies were added for 1.5 hours at RT. To detect HRP activity, Enhanced Chemiluminescence (ECL) Western blotting substrate (BioRad) was used. Western blot images from film developed using a SYNGENE G: Box imaging system (with GeneSnap software) were analyzed using Image J software [25].

### 2.6. Surface Plasmon Resonance (SPR)

SPR was performed using the BiaCore T100 (GE Healthcare) instrument to evaluate the binding affinity of protein molecules. The immobilization was done by activating carboxymethyl groups on a dextran-coated chip with N-hydroxysuccinimide. Subsequently, proteins were covalently bonded to the chip surface *via* amide linkages, followed by blocking these sites with ethanolamine to prevent any excess activated carboxyls. Reference surfaces were prepared similarly, except all carboxyls were blocked without adding protein. Each cell with an immobilized fusion polypeptide was paired with an adjacent reference cell on the chip during analysis. The concentration of bound protein was calculated by subtracting the reference RU from the protein RU. Phosphate-buffered saline with 5% DMSO served as both a running buffer and an analyte-binding buffer. A ligand molecule (analyte) flowed over the immobilized protein surface, and the binding response of the analyte to the protein was recorded. The maximum RU for each analyte indicated the interaction level and its relative binding affinity.

After normalizing the data along the x- and y-axes, the blank bulk refraction curves from the control flow chamber for each injected concentration were subtracted. The resulting binding curves were displayed, and the association (Ka) and dissociation (Kd) rate constants were determined using BIAevaluation 4.1 software with its equation for 1:1 Langmuir binding. These values were then used to calculate the binding affinities (K_D_).

### 2.7. Microscale Thermophoresis (MST)

The recombinant SIRT1 protein was labeled with NT-647 fluorescent dye, and its concentration was adjusted to 10 μM in 100 μl using the labeling buffer from NanoTemper. The fluorescent dye, dissolved in 100% DMSO, was added in a 2-fold molar excess and incubated for 30 minutes at room temperature. Excess dye was removed using a column equilibrated with 50 mM Tris pH 8.0, 150 mM NaCl, 10 mM MgCl2, and 0.05% Tween-20. The fluorescence intensity of the labeled protein was measured at 650 nm and 280 nm. A fixed amount of labeled protein was then mixed with 16 serial dilutions of purified N-terminal TULP3 (ligand) in a 1:1 ratio and loaded into the Monolith standard capillaries, and thermophoresis was measured on a Monolith NT.115 equipment (NanoTemper Technologies, Germany). The N-terminal TULP3 concentration range in the reaction mixture spanned from 320 μM to 10 nM. All the experiments were carried out in triplicate. The MO Affinity Analysis 2.2.7 software was used to analyze the acquired data. The MST run yielded the highest signal-to-noise ratio and lowest standard error for the K_D_ determination. The binding constants (K_D_) were calculated by fitting the data with a 1:1 stoichiometry according to the law of mass action.

### 2.8. Deacetylation activity of SIRT1 and SIRT2

In the deacetylation assay, a fluorescently labeled peptide, RHKK[ac]-Coumarin, based on the p53 sequence with acetylation at Lys382, was used as a substrate. The assay’s principle involves the deacetylation of the fluorescent peptide by Sirtuin in the presence of the co-substrate NAD+. The addition of trypsin then cleaves the fluorescent AMC-tag (7-amino-4-methylcoumarin), resulting in an increased fluorescent signal. The assay was performed by adding 250 µM of Sirtuin, 100 µM of the AMC peptide, and 500 µM NAD+ in the assay buffer (50 mM Tris pH 8.0, 137 mM NaCl, 2.7 mM KCl, 1 mM MgCl2, 1 mg/ml BSA), followed by a 30-minute incubation at 37°C. After incubation, a developer mixture containing 2 mM nicotinamide (NAM) and 10 mg/ml trypsin was added to the reaction mixture and incubated for 45 minutes at room temperature. Fluorescence measurements were conducted on a microplate reader (Infinite M200, TECAN) with an excitation wavelength of 360 nm and an emission wavelength of 460 nm. A reaction mixture containing all assay components except the enzyme served as a blank and was subtracted from the samples containing the enzyme.

### 2.9. Bioinformatics analysis

Nucleotide and protein sequence similarity searches, along with the identification of conserved domains, were performed using BLAST, CLUSTAL Omega, and CDD tools, available from the National Center for Biotechnology Information (https://blast.ncbi.nlm.nih.gov/Blast.cgi), the Conway Institute (https://clustal.org/omega), and (http://ccd.rhpc.nki.nl). Protein secondary structures, disordered regions, and domain predictions were analyzed using Alfa-fold, PSIPRED 4.0, DISOPRED3, and DomPRED (http://bioinf.cs.ucl.ac.uk/psipred/).

### 2.10. Statistical analysis

The data were collected from three independent experiments, and all values presented are expressed as the mean ± standard deviation. Statistical significance was evaluated with the ANOVA test in GraphPad Prism 8.0.1.

## 3. Results

### 3.1. Secondary and tertiary structure analysis of the Tubby-like proteins

Our study employed a structural genomics approach to investigate human Tub domain-containing proteins, focusing on TULP3. Multiple expression constructs were designed based on previously published crystal structures. TULP proteins are characterized by two domains: an N-terminal transcription activation domain and a C-terminal DNA binding tubby domain. The Tubby domain, spanning approximately 270 amino acids, exhibits high specificity in binding to both DNA and phosphatidylinositol. The tertiary structure of the Tubby domain in TULP features a β-barrel and an α-helix surrounding a central pore. The TULP3 gene is located on chromosomal band 12p13 (Uniprot Id: O75386). Binding to phosphatidylinositol allows Tubby to function downstream of cytoplasmic receptors, particularly G protein-coupled receptors (GPCRs). Upon reaching the nucleus, the amino-terminal domain of Tubby stimulates transcription. This intricate mechanism underscores the multifaceted roles of TULP3 in cellular processes.

TUB and TULP1-3 are closely related within the vertebrate family of Tubby-like proteins, while TULP4 is more distantly related. Alignment with the structure of TUB family proteins (Tub and TULP 1-3) reveals that all members share the conserved C-terminal tubby domain. However, the N-terminal region is not as conserved as the C-terminal, leading to the classification of tubby-like proteins. As shown in fig.1, the N-terminal region (amino acids 13-43) forms a conserved helix in the TULP3 model, crucial for binding to the IFT-A complex, and this helix is conserved across all four proteins in the tubby-like protein family. The tubby domain of TULP3 exhibits 70% sequence similarity with TUB and 66% with TULP1. The C-terminal domain is essential for stability and proper folding, ensuring functionality even in specific sub-cellular locations. The structure-based sequence alignment of all four tubby family proteins shows the conserved positively charged phosphate-coordinating residues, Lysine and Arginine (K268 and R270 in TULP3) residues involved in the plasma membrane localization. In contrast, TULP4 possesses a distinguishable WD40 domain, known for its role as a platform for protein-DNA or protein-protein interactions, and a SOCS (suppressor of cytokine signaling) box. While the full-length structure of TULP3 has not been solved, the N-terminal region was recently determined in a complex with IFT-A (PDB-8FH3). An Alpha Fold model, including the tubby domain, is also available for the TULP3 structure (Fig 2A). When aligned with TULP1 (2FIM) and TULP2 (1S31) structures, TULP3 demonstrates conservation of the tubby domain (Fig 2B). The structural analysis and sequence alignment of TULP3 indicate its potential dual localization (nucleus and plasma membrane) and a signaling role. As mentioned by Wang et al., TULP3 might function as a transcription factor in response to signaling [27]. This structural information enhances our understanding of the functional diversity within the Tubby-like protein family.

**Figure 1.**
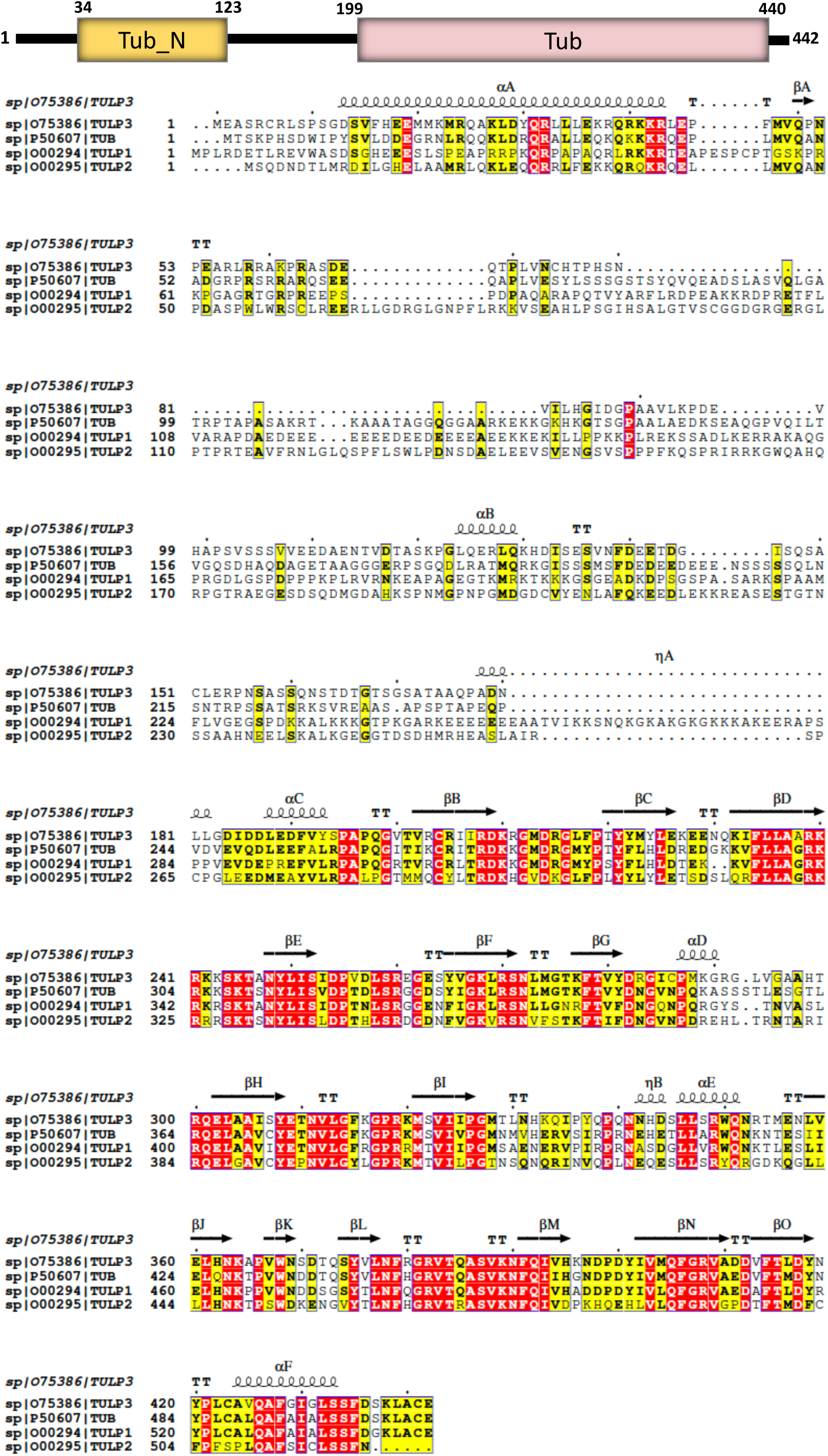
Structure-based sequence alignment illustrating sequence conservation among human Tubby-like proteins. All 4 family members are aligned to assess the domain sequence similarity. The secondary structure is indicated above the sequence. The blue box above the sequence indicates that the sequences of at least two out of four proteins share a similarity, while strictly conserved residues are shown in white on a red background. The sequence alignment was performed using ESPript 3.0 server [26]. The schematic representation of TULP3 domains is shown above the aligned sequence.

**Figure 2.**
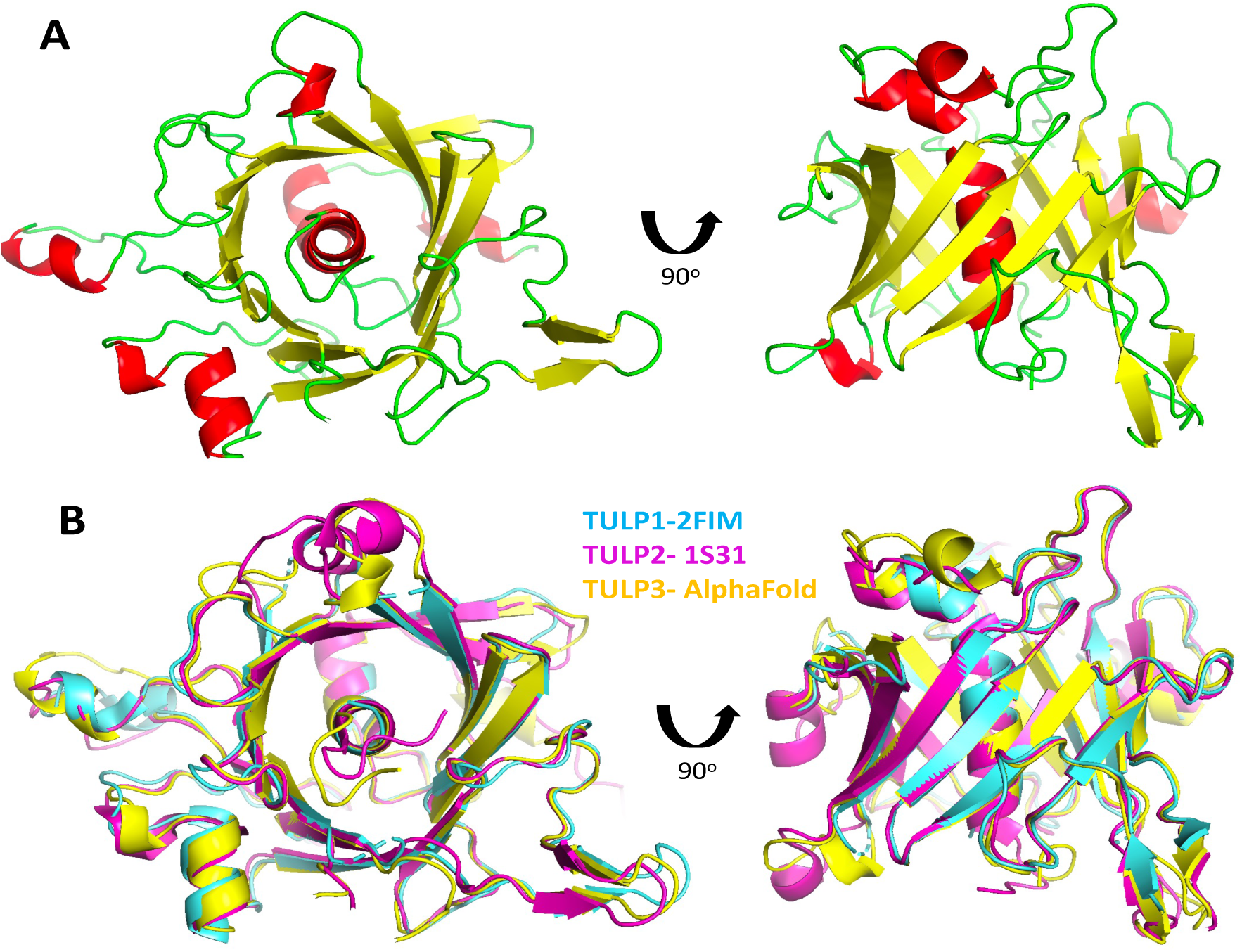
The AlfaFold homology model was built for C-terminal TULP3. (A) The top and front view of the tubby domain in the homology model from AlfaFoldDB. The tubby domain has a helix-filled β-barrel structure with 12 β-sheets and a hydrophobic helix that is filling the barrel. β-sheets and helices are depicted in yellow and red color, respectively. (B) Structure alignment of TULP3 model against crystal structure of TULP1 (PDB Id: 2FIM) and TULP2 (1S31).

### 3.2. TULP3 constructs and protein expression

In the expression and purification of TULP3, SIRT1, and SIRT2, specific constructs (Fig. 3) were designed for protein expression. Initial attempts with a C-terminal His-tag for TULP3 did not yield stable proteins, leading to the generation of new constructs with an N-terminal GST tag. Various purification techniques were employed to optimize the purification process. Since the isoelectric point (pI) of the protein from both Construct 1 and Construct 2 was the same, consistent buffers were utilized throughout the purification steps.

**Figure 3.**
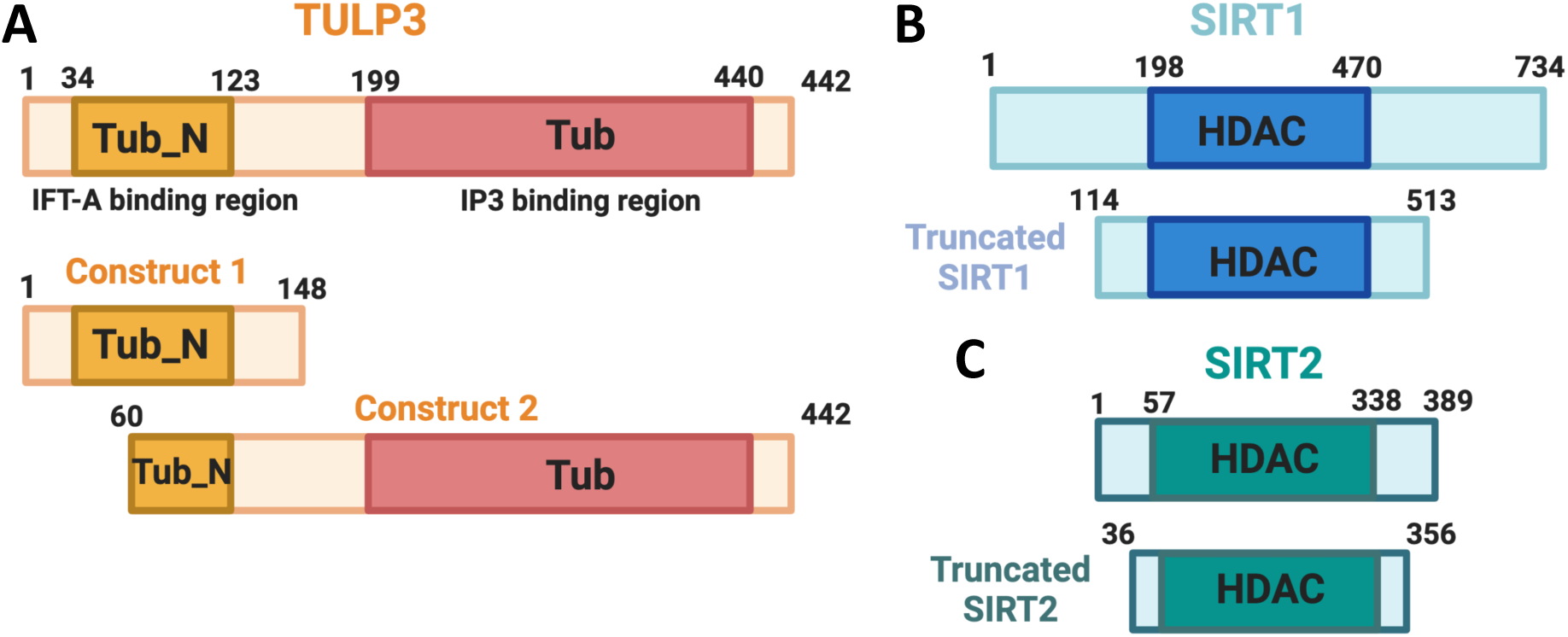
Construct preparation for the TULP3 and SIRT Proteins. The schematic representation of constructs used for the study; (A) N-terminal TULP3 (1-148 aa, Construct 1) and TULP3 (60-442 aa, Construct 2). (B) SIRT1 (114-513 aa). C) SIRT2 (36-356 aa).

The TULP3 proteins were expressed and purified as detailed in the materials and methods section. Protein stability was controlled using 10% glycerol, and it was observed that the protein was most stable at 2 to 5 mM DTT. The GST-affinity purified protein was then analyzed on SDS-PAGE, revealing the protein with GST at an expected molecular weight of ∼45 kDa and 69 kDa for Construct 1 and Construct 2, respectively. Subsequently, the dialyzed protein underwent cleavage to remove the GST-tag using precision protease. Through affinity chromatography, undigested protein and contaminants were removed, and the protein was further subjected to gel filtration to achieve high purity. During the final concentration step, buffer exchange was performed to replace DTT with 1 mM TCEP (tris(2-carboxyethyl)phosphine). Sirtuin proteins were purified using similar protocols as outlined in the materials and methods section. The purified proteins were concentrated to 1 mg/ml and then flash-frozen before storage at −80°C.

### 3.3. Protein characterization of TULP3

The concentrated TULP3 protein (construct-2, 8 - 10 mg/ml) with 1 mM TCEP exhibited stability upon adding Inositol phosphate-3 (IP3). The characterization of the protein was conducted through various methods:

1. Mass Spectrometry (Fig. 4A): Sequence analysis using mass spectrometry provided insights into the composition of the TULP3 protein. The analysis revealed peptides covering 44% of the complete protein sequence.
2. Size-Exclusion Chromatography (Fig. 4B): The SEC purification of N-terminal TULP3 (Construct-1) distinctly reveals the protein’s dimeric nature. Elution profiles were compared with those of known standardized proteins (Fig. 4B), facilitating the determination of TULP3’s size relative to the standards and confirming its dimeric state. This technique helps determine the protein’s oligomeric state and overall structural integrity.
3. Native PAGE and Western Blotting with Anti-TULP3 Antibody (Fig. 4C): TULP3 proteins (Constructs 1 and 2) were assessed for oligomerization using Native-PAGE. The N-terminal TULP3 (Construct 1) demonstrated dimerization around 35 kDa, while TULP3 (Construct-2) exhibited a band shift from 42 kDa, positioned above the 67 kDa BSA band (Fig. 4C). Similar dimeric patterns were noted in other tubby-like proteins, consistent with crystal structures showing homodimer in the asymmetric unit (e.g., TULP1, PDB: 2FIM, 3C5N). This implies a common oligomerization feature among tubby-like proteins. Western blotting using an anti-TULP3 antibody was performed to confirm the presence and identity of the TULP3 protein in the sample. It provides specificity in detecting the target protein, among other components. These analyses collectively contribute to comprehensively characterizing the TULP3 protein, confirming its stability, structural integrity, and presence in the sample.

**Figure 4.**
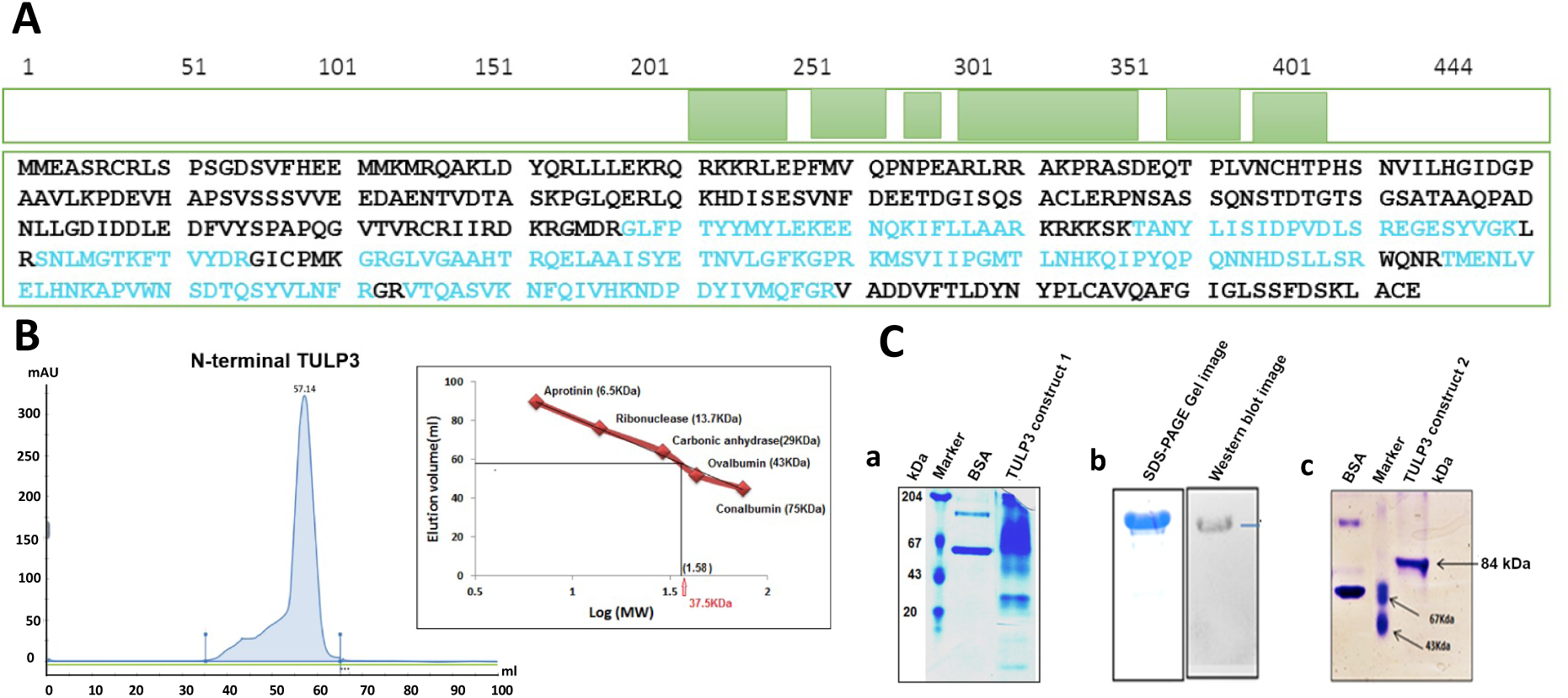
(A) Peptides analyzed from TULP3 protein (Construct-2). The peptide analysis showed 44 % coverage for the entire protein sequence. Peptides that are identified are marked in blue color. The coverage is shown in green marking. (B) Dimerization of N-terminal TULP3 protein (Construct-1). The chromatogram from the size-exclusion chromatography shows a peak for the N-terminal protein elution at 57.14 ml. The inner figure displays a modified calibration curve determined from five standard proteins (from Gel Filtration Calibration Kit LMW; GE Healthcare, USA) on Superdex 75 16/600 (GE Healthcare, USA): Conalbumin (75,000Da), Ovalbumin (43,000Da), Carbonic anhydrase (29,000Da), Ribonuclease A (13,700Da) and Aprotinin (6,500Da). (C) Protein characterization of TULP3 (Constructs 1 & 2) proteins. (a) Western blot for the N-terminal TULP3 protein (construct-1) displayed a band near 35 kDa. (b) TULP3 protein (Construct-2), where a C-terminal tubby domain-specific antibody, was used as the primary antibody. (c) Native gel for the TULP3 protein (Construct-2) presented a band above 67 kDa marker band that proves that the protein is a dimer.

### 3.4. The interaction between the HDAC domain of SIRT1 and SIRT2 with TULP3 constructs

The interaction between SIRT1, its activator AROS [28], and its inhibitor DBC1 [29,30] is well-established. However, the interaction of TULP3 with sirtuins has not been thoroughly studied. To address this gap, we conducted comprehensive studies to analyze the binding of TULP3 with SIRT1 and SIRT2 using pull-down assays (Fig. 5), surface plasmon resonance (SPR) experiments (Fig. 6), and microscale thermophoresis (MST) analysis (Fig. 7). These investigations aim to shed light on the molecular interactions between TULP3 and Sirtuins, providing valuable insights into their potential functional associations.

**Figure 5.**
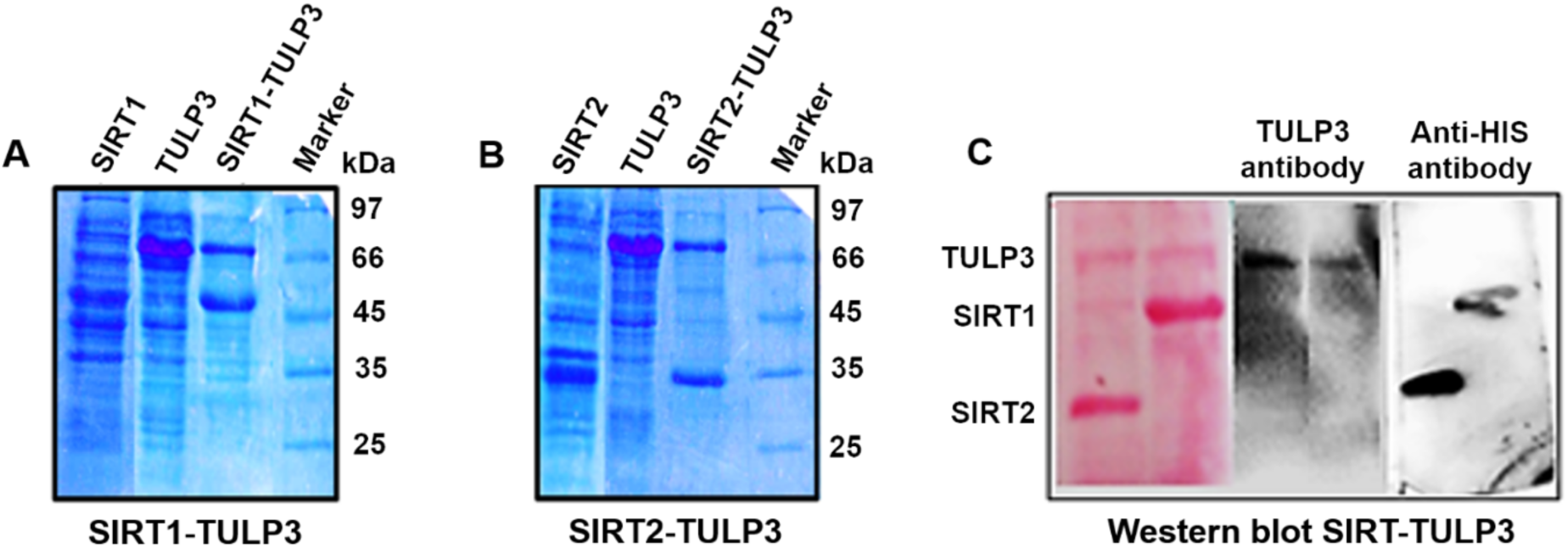
Pull-down assay of SIRT1 and SIRT2 with the TULP3 Construct 2. SDS-PAGE shows the bands for A) SIRT1 and TULP3, B) SIRT2 and TULP3 in the eluted sample after the pull-down experiment. SIRT1-TULP3 and SIRT2-TULP3 are the eluents of SIRT1 and SIRT2, respectively, in complex with TULP3 protein. His-SIRT1, His-SIRT2, and GST-TULP3 bands are at 46 kDa, 35 kDa, and 66 kDa, respectively. C) The pull-down complexes were then assessed by Western blot, where anti-His and anti-C-terminal TULP3 primary antibodies were used to check the presence of SIRT1/2 and TULP3, respectively.

**Figure 6.**
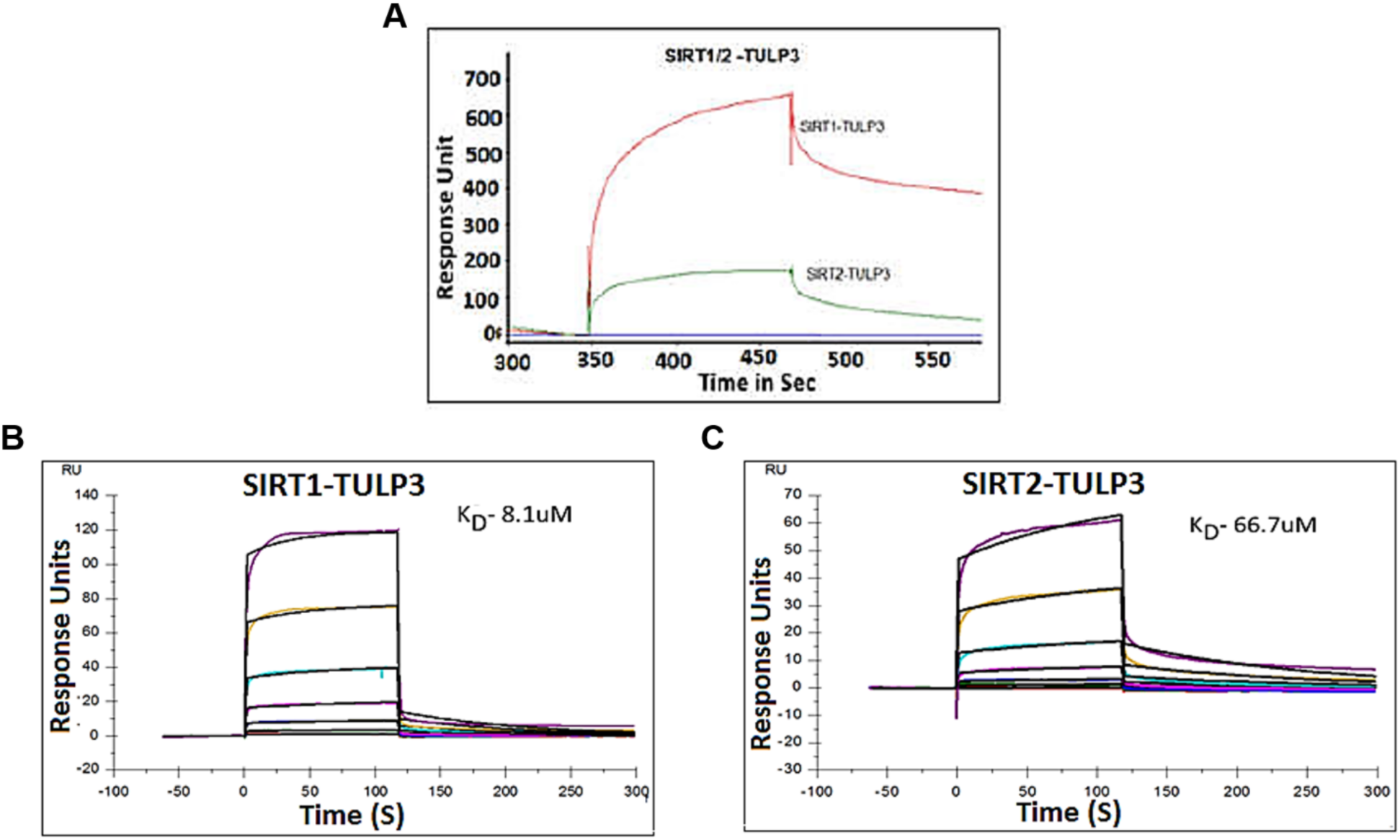
Quantitative binding studies for SIRT1 and SIRT2 interaction with TULP3 protein (construct-2) by SPR. A) Chromatogram showing binding of SIRT1-TULP3 and SIRT2-TULP3 complex. B) Sensogram for the titration of SIRT1 against TULP3 immobilized on a CM5 chip. The derived dissociation constant K_D_ (8 µM) was calculated from the rate constants. C) Sensogram of the titration of SIRT2 against TULP3. The derived dissociation constant K_D_ (66.7 µM) was calculated from the rate constants.

**Figure 7.**
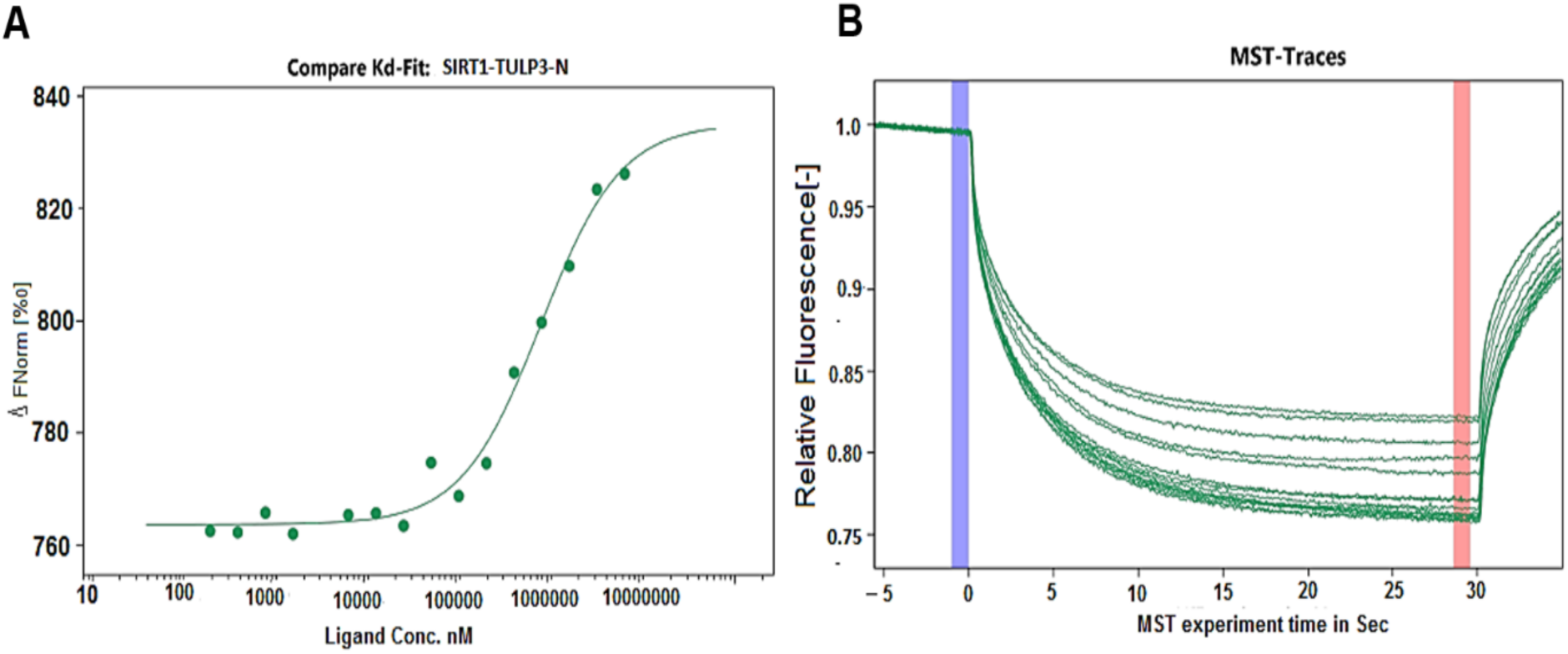
MST analysis of SIRT1 interaction with N-terminal TULP3 protein (construct-1). A) Binding curves were shown as the change in normalized fluorescence (ΔFnorm). B) The combined traces illustrate an MST experiment with 16 capillaries, each containing a constant concentration of the fluorescent interaction partner and varying concentrations of the non-fluorescent partner. All traces are normalized to an initial relative fluorescence value of 1. F_Norm_ values at a specific time point were plotted against the inhibitor concentration, and the Kd was determined by fitting the data to a hyperbolic binding curve.

#### 3.4.1. The TULP3 (Construct-2) interaction with SIRT1 and SIRT2 by pull down assay

SIRT1 and SIRT2 constructs, incorporating the HDAC domain, were employed for both pull-down and SPR analyses. In the pull-down assay, His-SIRT1/SIRT2 was immobilized on Ni-NTA beads, followed by thorough washing with TBS buffer to eliminate unbound proteins. Subsequently, TULP3 was introduced to bind to the bait in the column. The elution of the SIRT1-TULP3 and SIRT2-TULP3 complexes was confirmed by SDS-PAGE. A subsequent Western blot, using an anti-His antibody and anti-TULP3 antibody, verified the presence of bait and prey in the complexes (Fig 5). These experiments provide robust evidence of the interaction between TULP3 and both SIRT1 and SIRT2.

#### 3.4.2. The TULP3 (Construct 2) interaction with SIRT1 and SIRT2 by SPR

The SDS-PAGE and Western blot results unambiguously demonstrate the interaction between TULP3 and both SIRT1 and SIRT2. An SPR analysis was conducted to confirm this interaction further. SIRT1 and SIRT2 were immobilized on a CM5 chip using 1X PBS buffer after pH scouting. The purified TULP3 (Construct 2) protein was then used to check binding with SIRT1 and SIRT2. The SPR binding study revealed that TULP3 exhibits a stronger interaction with SIRT1 than SIRT2 (Fig 6A). Subsequent kinetic analysis was performed to determine the K_D_ value. Different concentrations of SIRT1 and SIRT2 proteins were passed over the immobilized TULP3 on the chip, and the K_D_ values were deduced using the 1:1 Langmuir binding equation. For SIRT1-TULP3 and SIRT2-TULP3 interactions, K_D_ values were found to be 8.1 µM and 66.7 µM, respectively (Fig. 6B and C). This further supports the notion that, as observed in the pull-down assay, TULP3 has a stronger interaction with SIRT1 than SIRT2.

#### 3.4.3. The N-terminal region of TULP3 (Construct 1) interacts with SIRT1 by MST

To identify the binding region of TULP3 with HDAC domain, both the TULP3 constructs were used. Both the TULP3 constructs with GST tag showed the binding with SIRT1 and SIRT2 as seen in the binding analysis by SPR. N-terminal TULP3 showed significant binding affinity (K_D_) of 28 µM with SIRT1 (Fig 7). Since the TULP3 (Construct 2) has a small portion of the Tub N domain, we speculate that the Tub N domain of TULP3 is likely to interact with the HDAC domain of SIRT1. The assessment of the C-terminal tubby domain binding interaction with SIRT1 would give us better information about the region of interaction of the TULP3 protein. In our study, since the C-terminal protein was insoluble, we could not use it for the binding studies.

### 3.5. The effects of TULP3 on SIRT1 deacetylation activity

The deacetylation assay was conducted using pure TULP3 proteins and SIRT1 cell lysate. Serially diluted TULP3 construct proteins were titrated against SIRT1, revealing no significant change in the activity of SIRT1 across all concentrations of TULP3 constructs (Fig. 8A and B). In contrast, resveratrol at a concentration of 100 µM exhibited a >50% increase (p<0.0001) in SIRT1 deacetylation activity. The Cayman SIRT1 direct fluorescent assay kit was employed for the titration. Given that TULP3 construct proteins do not impact the activity of SIRT1, it is speculated that TULP3 neither acts as an activator nor an inhibitor of SIRT1 deacetylase activity. This raises the question of whether TULP3 itself serves as a substrate for SIRT1. The TULP3 protein sequence was analyzed for potential acetylated lysine sites (Table 1) using a webserver (ASEB; A Web Server for KAT-specific Acetylation Site Prediction) and acetylation set enrichment-based method [31]. Several potential SIRT1-specific lysine residues were identified, suggesting TULP3 as a substrate for SIRT1. However, a recent study demonstrated that TULP3 is deacetylated by HDAC1, not by SIRT1 [15].

**Figure 8.**
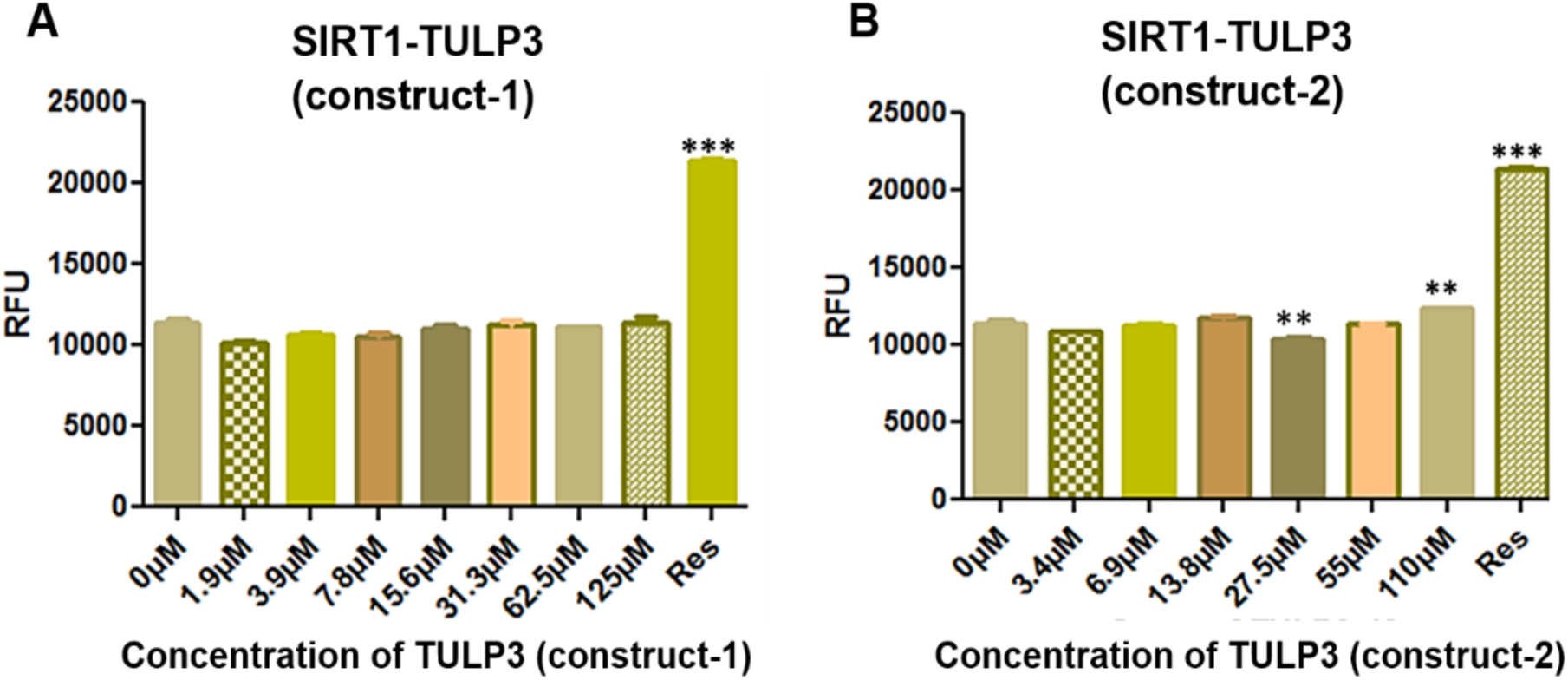
Deacetylation activity of SIRT1 in the presence of TULP3 proteins. The deacetylation assay was performed by titrating serially diluted TULP3 proteins. The experiment was performed by using Cayman SIRT1-direct deacetylation kit by taking, A) 0-125 µM of N-terminal TULP3 (construct-1) and B) 0-110 µM of the TULP3 (construct-2) proteins. No change was observed in the activity of SIRT1 with the TULP3 construct proteins. 250 µM concentration of resveratrol was used as a control.

**Table 1.**
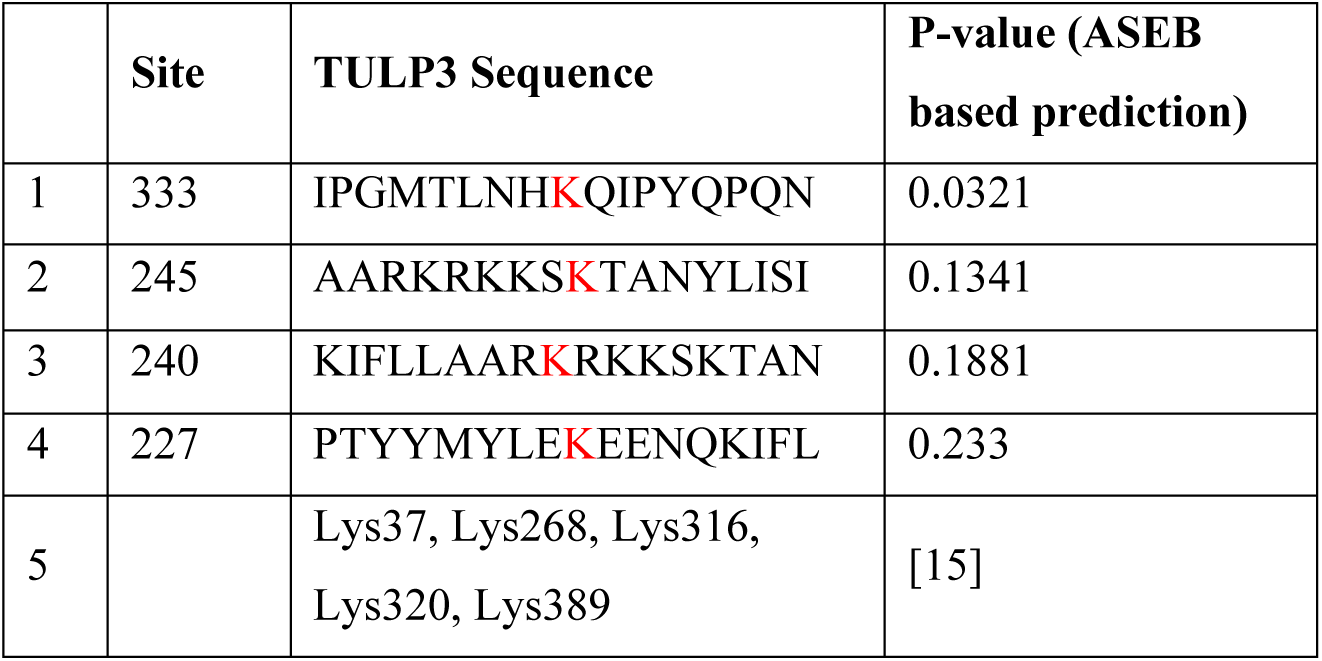
Acetylation sites in TULP3.

## 4. Discussion

Tubby-like protein 3 (TULP3) is part of the tubby family, involved in neural cell functioning and maintenance [7,32]. Tubby family proteins, including Tub and Tulps1-3, facilitate the ciliary trafficking of membrane proteins like G protein-coupled receptors and polycystin 1/2 complex [13,33]. TULP3 has been implicated in embryonic neural tube development and is also involved in the Hedgehog signaling pathway [32]. Mutations in the parent Tub gene can lead to adult-onset obesity, retinal and cochlear degeneration in mice, and abnormalities such as exencephaly, spina bifida, and facial clefting [3,6,34]. Sirtuins regulate various physiological processes including cellular differentiation, apoptosis, metabolism, aging, immune response, oxidative stress, and mitochondrial function. Their deacetylation activity is crucial for epigenetic alterations implicated in autoimmune diseases. SIRT1 and SIRT2, both NAD-dependent protein deacetylases, play protective roles against age-related diseases such as cancer, diabetes, cardiovascular disorders, neurodegenerative diseases, and cellular senescence [35–37]. The involvement of Tubby family proteins in these cellular processes underscores their significance in regulating cellular signaling and sensory functions, emphasizing the importance of understanding their cellular interacting partners.

Structure-based sequence alignment showed the N-terminal region is conserved in all the 4 Tulp proteins; it consists of α–helix, and also C-terminal tubby domain is highly conserved between 4 Tulp’s consisting of α –helix with β–barrel sheets (Fig 2). It shows that the C-terminal tubby domain is more conserved as compared with the N-terminal domain.

The characterization of TULP3 protein was validated by size-exclusion chromatography and Native-PAGE, which clearly showed that TULP3 is a dimer (Fig 5B). We can see the homologous crystal structure of TULP1 as a dimer, where two monomers have been seen in the asymmetric unit (PDB: 2FIM, 3C5N). Further, validates that TULP3 protein is a dimer. Peptide analysis of TULP3 protein showed 44 % coverage for its sequence by mass spectrophotometry. Western blot was performed for the TULP3 Constructs 1 and 2 proteins which were determined by using specific antibodies.

A proteomic approach was undertaken to identify crucial interacting partners of SIRT1, shedding light on its involvement in regulating physiological and pathological processes. Mass spectrometry (LC-MS/MS) was employed to uncover these interacting partners, revealing TULP3 as one of them. TULP3, known for its role in signal transduction and transcription, emerged as a potential interacting partner of SIRT1 [38]. The initial step to confirm the interaction between SIRT1, SIRT2, and TULP3 involved pull-down experiments. This evidence supports the notion that there is a physical interaction between the sirtuins (SIRT1 and SIRT2) and TULP3, laying the foundation for further exploration into the nature and significance of this interaction. Continuing the investigation, biophysical techniques, specifically surface plasmon resonance (SPR), were employed to further validate the interaction between TULP3 and SIRT1. The SPR results revealed that SIRT1 interacts with TULP3, showing a K_D_ value of 8.1 μM. In contrast, SIRT2 exhibited a lower binding efficiency with a K_D_ value of 66.5 μM. Moreover, the Microscale Thermophoresis assay results provided conclusive evidence of a robust binding association between the N-terminal of TULP3 (Construct 1) and the SIRT1 protein. These biophysical confirmations strengthen the evidence that TULP3 is a bona fide *in vivo* partner for SIRT1/2, contributing to a more comprehensive understanding of their interaction dynamics.

The acetylation of specific residues, Lys316 and Lys389, on TULP3 by p300 has been identified as a regulatory mechanism that enhances the stability of TULP3 protein. Conversely, HDAC1 counteracts this effect by deacetylating these lysine residues. This deacetylation process mediated by HDAC1 leads to decreased TULP3 protein levels, primarily through the cullin3 RING ligase complex and subsequent proteasomal degradation pathway. The intricate interplay among p300, HDAC1, and cullin3 proteins highlights an acetylation switch that finely tunes TULP3 protein stability [15]. This study supports the role of HDAC1 as a deacetylase for TULP3 protein, contributing to our understanding of the post-translational regulation of TULP3.

A deacetylation experiment was conducted to investigate the molecular interactions and functions of TULP3 in relation to SIRT1/2. The assay revealed no significant change in the deacetylase activity of SIRT1 and SIRT2 in the presence of TULP3. It suggests that TULP3 may not be an activator or inhibitor of sirtuin deacetylase activity, in contrast to the previously suggested idea of TULP3 being a substrate for sirtuin [15]. Hence, we speculate that TULP3 might have other cellular functions upon binding to sirtuin. The family member, TULP1, is known to be involved in DNA repair that interacts with SIRT1.

The study investigated the interaction between TULP3 and SIRT1/SIRT2 proteins using various biophysical and biochemical methods. Substantial binding interactions were observed between TULP3 constructs and SIRT1/SIRT2. Contrary to expectations and previous assumptions, the results indicated that TULP3 does not act as a substrate for SIRT1/SIRT2. This opens up new avenues for exploring the cellular functions and interactions involving TULP3 and SIRT1/SIRT2.

## Abbreviation

TULP3: Tubby-like protein 3
Sirtuin: Silent mating type information regulation 2 homolog
GPCRs: G protein-coupled receptors
IFT-A: Intraflagellar transport - A
HDACs: Histone deacetylases
GST: Glutathione S transferase
PIP2: Phosphatidylinositol 4, 5-bisphosphate
NF-kB: Nuclear factor kappa B
IPTG: Iso propyl thio-β-galactoside
PMSF: Phenyl methyl sulfonyl fluoride
EDTA: Ethylene diamine tetra acetic acid
Ni-NTA: Nickel Nitrilotri acetic acid
TBS: Tris buffered saline
SDS-PAGE: Sodium dodecyl sulfate polyacrylamide gel electrophoresis
SPR: Surface plasmon resonance
DMSO: Dimethyl sulfoxide
MST: Microscale thermophoresis
NT-647: Nanotemper-647
NAD: Nicotinamide adenine dinucleotide
AMC: 7-amino-4-methyl coumarin
NAM: Nicotinamide
IP3: Inositol phosphate-3
TCEP: Tris (2-carboxyethyl) phosphine
DBC1: Deleted in bladder cancer protein 1
AROS: Active regulator of SIRT1
CM5: Carboxy methylated 5
PVDF: Polyvinylidene difluoride
PDB: protein data bank

## Author contributions

BP and RM designed the research; RM and GK performed molecular biology, protein production, and biochemical assays. VS performed bioinformatics analysis. BP, RM, and GK wrote the original draft; all authors contributed to the final version of the manuscript.

## Acknowledgments

BP is grateful to the Department of Biotechnology (DBT), the Government of India for the financial support (BT/PR32378/BRB/10/1784/2019), and the Department of Science & Technology (DST), Government of India (DST-FIST: SR/FST/LS-I/2017(C)). MR is grateful to the Council of Scientific and Industrial Research (CSIR) for the SRF fellowship (09/490(0093)2012-EMR-1). GK is grateful to the Indian Council of Medical Research (ICMR), Government of India, for the SRF fellowship (45/38/2019-BIO/BMS).

## Notes

### Competing Interest Statement

The authors have declared no competing interest.

